# seqgra: Principled Selection of Neural Network Architectures for Genomics Prediction Tasks

**DOI:** 10.1101/2021.06.14.448415

**Authors:** Konstantin Krismer, Jennifer Hammelman, David K. Gifford

## Abstract

Sequence models based on deep neural networks have achieved state-of-the-art performance on regulatory genomics prediction tasks, such as chromatin accessibility and transcription factor binding. But despite their high accuracy, their contributions to a mechanistic understanding of the biology of regulatory elements is often hindered by the complexity of the predictive model and thus poor interpretability of its decision boundaries. To address this, we introduce seqgra, a deep learning pipeline that incorporates the rule-based simulation of biological sequence data and the training and evaluation of models, whose decision boundaries mirror the rules from the simulation process. The method can be used to (1) generate data under the assumption of a hypothesized model of genome regulation, (2) identify neural network architectures capable of recovering the rules of said model, and (3) analyze a model’s predictive performance as a function of training set size, noise level, and the complexity of the rules behind the simulated data.

## I. Introduction

Over the last five to ten years, neural networks were successfully applied to make large gains on a wide range of tasks in such diverse fields as computer vision, computer audition, natural language processing, and robotics. While the structure and the semantics of the data used to train and evaluate neural networks can be vastly different, the core learning algorithms are almost always the same and the neural network architectures are often composed of similar building blocks. This is also true for the field of genomics, and computational biology as a whole, where deep neural networks are trained on data that are obtained experimentally using functional genomics assays such as DNase-seq [1], ATAC-seq [2], and ChIP-seq. Motivated by their success, architectural building blocks commonly seen in these networks, such as convolutional layers, recurrent layers, batch normalization, drop-out, and skip connections [3–6], have been imported from computer vision and other fields. This cross-fertilization between fields and the general applicability of the building blocks of deep learning has more recently been seen in the adoption of transformer-based architectures for image classification tasks in computer vision and protein prediction tasks in biology. However, most data sets used to train supervised deep learning models in biology are different from data sets in computer vision and natural language processing in two ways. (1) Biological problems contain noisy input and noisy labels in that not only is there substantial intra-class variability and noise in the input, e.g., images labeled as *cat* contain cats that vary in terms of breed, color, position, pose, etc., but also a significant fraction of examples are mislabeled, i.e., images labeled as *cat* are empty or contain dogs. This is rare in computer vision data sets, but common in data sets derived from functional genomics assays. (2) Feature attribution or other model explanation methods are not human-interpretable. We understand images of cats in the sense that we know which parts of the image contain information that is relevant for the classification (because they belong to the cat) and which parts are irrelevant (because they belong to the background). This intuitive understanding is necessary when attribution methods such as saliency maps are applied to assess a model’s ability to base predictions on relevant parts of the input. In biology, examples often include DNA sequence windows of various widths, most commonly 1000 base pairs (bp), which, unlike images of cats, are not *human-readable*. This biology-specific issue of inherently opaque examples exacerbates the general interpretability issue of deep neural networks, whereas the lack of high quality data sets contributes to the reproducibility crisis and makes it more difficult to compare architectures, as they are often only evaluated on a custom data set.

The method introduced here, *seqgra*, attempts to improve the process by which neural network architectures are chosen for specific genomics prediction tasks and provides a framework to evaluate model interpretation methods. Its fully reproducible pipeline provides a means to (1) simulate data based on a pre-defined set of probabilistic rules, (2) create and train models based on a precise description of their architecture, loss, optimizer, and training process, and (3) evaluate the trained models using conventional test set metrics as well as an array of feature attribution methods. These feature attribution methods in combination with simulated data and thus perfect ground truth enable an analysis of the model’s decision boundaries and how well they capture the underlying rules of the data generation process from step 1. Utilizing this framework, models are not only evaluated based on their predictive performance, but also on the ability to recover the vocabulary (e.g., specific transcription factor binding site motifs) and grammar (e.g., spacing constraints between interacting transcription factors) of the data set, while assigning little weight to confounding factors and idiosyncratic noise.

Efforts in this area include Kipoi [7], a repository for trained genomics models, and Selene [8], a framework for biological sequence based deep learning models that supports training of PyTorch models, model evaluation with conventional test set metrics (ROC and precision-recall curves), and variant effect prediction and *in silico* mutagenesis of trained models. To our knowledge none of the existing methods offer functionality for simulating data using a general framework of probabilistic rules, nor do they incorporate feature attribution methods.

Furthermore, this simulation-based framework can also serve as a testbed for hypotheses about biological phenomena or as a means to investigate the strengths and weaknesses of various feature attribution methods across different neural network architectures that are trained on data sets with varying degrees of complexity. In the former use case, the hypothesis is encoded in the rules of the simulation process to identify an appropriate neural network architecture, which is subsequently trained and evaluated on experimental data. The performance of this simulation-vetted architecture on experimental data serves as an indication of the validity of the hypothesis and its underlying assumptions about the biological phenomenon.

## II. Materials and Methods

### A. Position probability matrices and position weight matrices

We use position probability matrices (PPM) with a DNA alphabet (Σ = {A, C, G, T}) to represent sequence motifs:

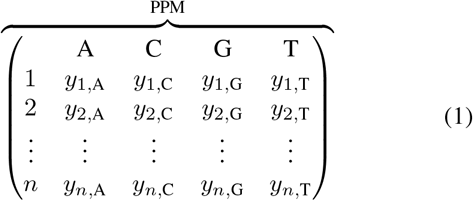

As the name suggests, each cell of a PPM is a probability, the probability of observing a particular nucleotide at a particular position, and each row sums to one, i.e., at each position one of the four nucleotides must be present. We use the notation PPM_*k*_(*i, j*) to access the probability of observing the *j*th nucleotide at the *i*th position in a specific PPM_*k*_.

These PPMs usually describe experimentally obtained estimates of transcription factor binding sites, but may also describe artificially constructed sequence motifs.

To calculate the likelihood of a sequence given a PPM, we first convert the PPM to a position weight matrix (PWM) by transforming the elements of the PPM to log likelihoods,

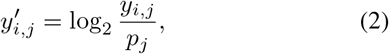

using background sequence probabilities *p*, which are described in section II-F. The *score* of a particular position in a DNA sequence is then calculated by adding the value of the observed nucleotide at each position in the PWM.

### B. Motif information content

To calculate the information content of a sequence motif represented as a PPM, we first calculate U(*i*), the uncertainty at position *i* as follows:

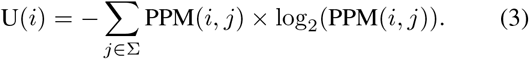

The information content at position *i* is then defined as follows

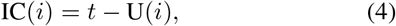

where *t* = log_2_(|Σ|), the total information content per position in bits. In order to obtain MIC, the information content of the entire motif, we add up the individual positions:

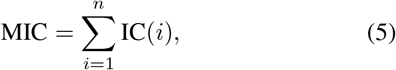

where *n* is the motif width in nucleotides (nt), see matrix in 1.

### C. Relative entropy between motif and background distribution

The information content of a motif is a special case of the relative entropy of a motif where background probabilities *p* are uniform. Relative entropy, also known as KL divergence, between a motif and the background distribution is calculated per position, similarly to IC:

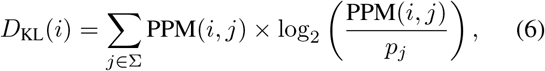

and then summed over positions to obtain the Motif Relative Entropy,

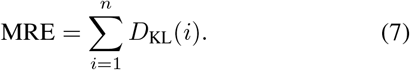

### D. Relative entropy between two motifs

While the relative entropy between a particular motif, PPM_1_, and the background distribution is a way to gauge the learnability of a grammar where the presence of PPM_1_ carries information, the relative entropy between two motifs, PPM_1_ and PPM_2_, is equally useful to assess the learnability of grammars with multiple, semantically distinct sequence elements.

By slightly adjusting the *D*_KL_ from above, we calculate the KL divergence of position *i* between two motifs as follows:

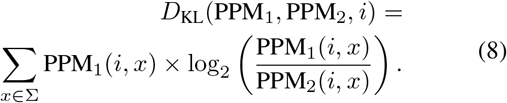

The motif pair relative entropy of PPM_1_ relative to PPM_2_ is then defined as

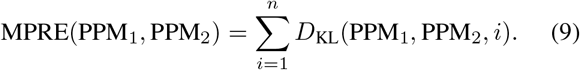

To calculate the MPRE between motifs of unequal width, we pad the shorter motifs with *neutral* positions using background probabilities.

Another issue with equation 9 is that it does not capture highly similar but shifted motifs. PPM_1_ might be equivalent to PPM_2_ shifted by one position and thus considered highly similar, but MPRE(PPM_1_, PPM_2_) in its current form does not reflect this. To resolve this, we calculate MPRE(PPM_1_, PPM_2_) for several alignments of PPM_1_ and PPM_2_ and take the minimum.

### E. Empirical similarity score between two motifs

The empirical similarity score (ESS) between PPM_1_ and PPM_2_ is another way to assess the similarity between two motifs and thus the difficulty to distinguish between them. ESS(PPM_1_, PPM_2_) is calculated by generating *k* (in this work, *k* = 100) instances of motif 2, flanked on both sides by background sequences of length *n*_1_, where *n*_1_ is the width of PPM_1_. All positions of these *k* sequences are then scored by PWM_1_ (the position weight matrix of PPM_1_), and the highest score per sequence is returned. ESS(PPM_1_, PPM_2_) is then the mean of these *k* scores. ESS motif matrix plots (Figure 2 and Supplementary Figure S3) depict adjusted empirical similarity scores, which are shifted by ESS_0_ if ESS_0_ < 0, where ESS_0_ = min_*j*_ ESS(PPM_*i*_, PPM_*j*_), and normalized such that the self similarity score ESS(PPM_*i*_, PPM_*i*_) = 1.0.

Both MPRE and ESS are asymmetric, i.e., ESS(PPM_1_, PPM_2_) ≠ ESS(PPM_2_, PPM_1_).

### F. Alphabet distribution for grammars

For all grammars discussed in this paper, we used the natural nucleotide distribution of the human genome, 29.565 % adenine (A), 20.435 % cytosine (C), 20.435 % guanine (G), and 29.565 % thymine (T) [9].

### G. Motif database

We used HOMER motifs for all grammar sequence elements that were based on transcription factor binding site motifs. These motifs were obtained by analyzing data from publicly available ChIP-seq experiments [10].

### H. Feature importance evaluators

While conventional test set metrics, such as ROC curves and precision-recall curves, assess model performance based on a set of examples (e.g., the test set), feature importance evaluators quantify the contribution of each input feature to the model’s prediction. In the context of seqgra, feature importance evaluators are used to assess what we call grammar or vocabulary recovery, the degree to which a model was able to align its decision boundaries with the rules of the grammar that was used to simulate the data it was trained on. This is possible because for simulated data we not only know the ground truth label for each example, but also which positions are part of the background and thus contain no information about the class label, and which positions were altered by a grammar rule and thus do contain information about the class label. These position-level annotations (*background positions*, *grammar positions*) are provided for all simulated examples.

More formally, feature importance evaluators take a model *f*(*x*), a target *y* and an example *x_i_* of width *n*, and return *z*, an *n*-dimensional vector that contains the attribution value (also known as importance, relevance, contribution) of each input position to model *f*(*x*) predicting target *y*. Please note that n is the sequence length of the example, not the number of features. For instance, if the input to the model is a 150 nt DNA sequence, *x_i_* is a 150 by 4 matrix (one-hot encoded), containing 600 features, but its width *n* = 150. Feature attribution values in seqgra are grouped and reported at the position level, not the input feature level.

Attribution values are visualized with so-called grammar agreement plots, which are heatmaps depicting attributions and position-level annotations of several examples. The plots encode the attribution values in the color luminosity, where lighter colors indicate low values (low feature importance) and dark colors indicate high values (high feature importance). The position-level annotations are encoded in the color hue, with grammar positions in green and background positions in red.

### I. Gradient-based feature importance evaluators

This large class of feature importance evaluators (FIEs) uses backpropagation to calculate the partial derivatives of the output, *f_y_*(*x*), with respect to the input, *x_i_*. seqgra includes seven gradient-based feature importance evaluators off-the-shelf, whose implementations are based on code by Yulong Wang [11].

The most basic FIE, **raw gradient** [12], just returns the gradient with respect to the input example *x_i_*:

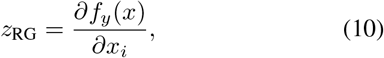

or short ∇*f_y_*(*x_i_*), where *f_j_*(·) is the activation of the target neuron in the output layer, e.g., class *j* for multi-class classification tasks.

The absolute gradient method or **saliency** is defined as

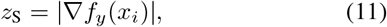

where |*x*| applies the element-wise absolute value operation to vector *x*.

**Gradient-x-input** [13] (gradient times input) is defined as

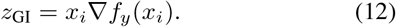

**Integrated Gradients** [14] takes the average of multiple (here, *K* = 100) gradients evaluated along the linear path from the baseline *x*_0_ (which in seqgra is the zero vector) to the input example *x_i_*. The method is defined as

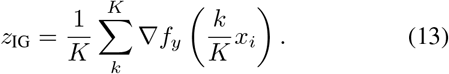

seqgra also supports gradient-based methods that alter the way the gradient is obtained using backpropagation, namely **Guided Backpropagation** [15], **Deconvolution** [16], and **DeepLIFT** [17]. The details of these methods are beyond the scope of this work.

### J. Model-agnostic feature importance evaluators

Model-agnostic FIEs do not require access to the gradients and make no assumptions about the structure of the model, hence the name. They rely solely on the ability to evaluate *f_y_*(*x*), for various altered versions of *x*.

**Sufficient Input Subsets** (SIS) [18] is a perturbation-based method that identifies subsets of input features that are sufficient to keep *f_y_*(*x*) > *τ*, i.e., if all other features are masked, the class prediction does not change (is still above some threshold *τ*). Unlike gradient-based FIEs, which return a real-valued vector of feature attributions, SIS returns a binary vector, indicating for each feature whether it is part of a sufficient input subset or not.

### K. Hardware infrastructure

Models presented in this paper were trained on three compute nodes with a total of 6 CPUs (2x Intel Xeon E5-2630 v4, 2x Intel Xeon Gold 6138, 2x Intel Xeon Gold 6240), 26 GPUs (8x NVIDIA GeForce GTX 1080 Ti with 11 GB GDDR5X, 10x NVIDIA GeForce RTX 2080 Ti with 11 GB GDDR6, and 8x NVIDIA Titan RTX with 24 GB GDDR6), and a total of 833 GB of main memory. The total GPU time (for training and evaluation) was roughly 12 GPU months.

### L. Software infrastructure

All seqgra data presented in this paper was obtained on machines running Ubuntu 18.04.3 LTS, CUDA 10.1, cuDNN 7.6.5, Python 3.8, NumPy 1.19.2, TensorFlow 2.2.0, PyTorch 1.7.0, and R 4.0.

## III. Results

### A. seqgra provides a reproducible, simulation-based framework for neural network architecture evaluation

The method we describe in this paper (seqgra) generates synthetic biological sequence data according to predefined probabilistic rules in order to either (1) evaluate neural network architectures trained on these data sets, or (2) test whether the assumptions about the underlying biological phenomenon that the probabilistic rules of the simulation process are based on, accurately reflect experimentally obtained data. In the former scenario, the result would be a neural network architecture that—when trained on data sets generated from a similar set of rules—has high predictive performance and decision boundaries that closely reflect those set of generative rules. The goal of the latter approach is to arrive at a concise set of probabilistic rules that approximates the biological process in question, and a neural network architecture whose high performance on simulated data is recapitulated when trained on experimental data.

A data set in the context of seqgra, whether obtained by simulation or experiment, is always divided into three subsets, training set, validation set, and test set. Each of the subsets comprises a number of supervised examples, which are (*x, y, a*)-triplets. Here, the input variable *x* is a biological sequence (DNA, RNA, protein) of fixed or variable length, also referred to as sequence window or features; *y* is the target variable, the *condition* this example belongs to (e.g., cell type), which is either a mutually exclusive *class* or a non-mutually exclusive *label*, for multi-class classification tasks or multi-label classification tasks, respectively; and *a* is the positional annotation of the example, denoting for each position in *x* whether it is part of the *grammar* or part of the *background*. Grammar positions contain information related to *y* and are therefore important for classification, whereas background positions do not and are thus irrelevant for classification.

The core functionality of seqgra can be broken down into three components: (1) Simulator, (2) Learner, and (3) Evaluator. Each component corresponds to a distinct step in the pipeline depicted in Figure 1A.

**Fig. 1.**
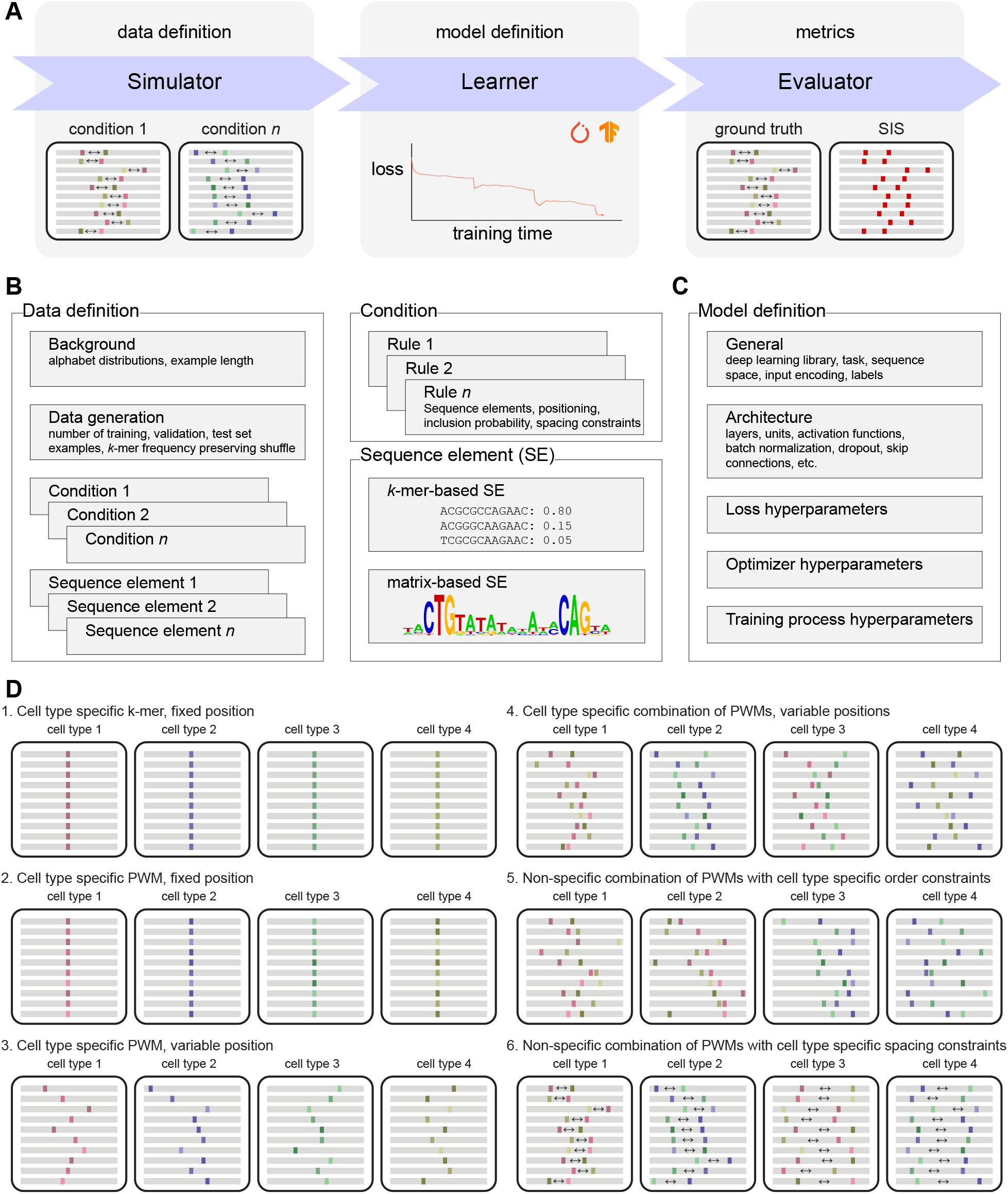
A framework for simulation-based evaluation of neural network architectures. (**A**) Schematic of the three main components: First, a simulator generates synthetic data according to the rules and specifications defined in the data definition file. Second, a learner creates a neural network model whose architecture and hyperparameters are specified in the model definition file, and trains it on the synthetic data from step 1. And third, the trained model is evaluated in terms of predictive performance and its ability to recover the rules specified in the data definition file. (**B**) The data definition specifies the basic properties of the synthetic data, including the alphabet (e.g., DNA, RNA, protein) and its distribution, as well as condition-specific rules (the *grammar*), which determine how information about the label *y* is encoded in the input *x*. (**C**) The model definition contains all information required to create and train the model. (**D**) A schematic of six simulated toy data sets for multi-class classification, where the classes *y* correspond to cell types and the input *x* are sequence windows (depicted as gray bars) that encode information about the class *y* at certain positions in *x* (colored areas). The rules that determine how this information is encoded range from basic (cell type specific *k*-mer at fixed position) to complex (non-specific combinations of position weight matrices with cell type specific spacing constraints).

In step 1, the simulator generates a synthetic data set according to the specifications laid out in the data definition (see Figure 1B), a document that contains a precise description of the generated data, from the background nucleotide distribution to the set of probabilistic rules that determines how information about the condition *y* (label, class) is encoded in the sequence window *x*. This set of probabilistic rules is also referred to as *grammar* or sequence grammar throughout this manuscript (hence the name *seq*-*gra*), and although related to formal grammars, seqgra’s probabilistic rules are not expressed as and not equivalent to production rules in the context of formal language theory.

Schematic depictions of six toy data sets, generated from probabilistic rules of varying complexity, are shown in Figure 1D. In each case, the data set contains examples belonging to one of four classes and the probabilistic rules determine how information about the class *y* (in this case, the cell type) is encoded in the sequence window *x*. The ability to recover this relationship during training is imperative for the model’s predictive performance. The sequence windows of the examples are shown as gray bars with colored spots, where background positions are shown in gray and grammar positions are shown in color. In the first example, each of the four cell types can easily be identified by the presence of a class-specific *k*-mer at the center of the sequence window, a relationship that, unsurprisingly, can be learned perfectly (i.e., close to an ROC AUC of 1.0) and efficiently (i.e., with few training examples) by most neural network architectures. Since a set of rules as simple as the one used in example 1 will almost always be an inadequate description of any biological process, seqgra allows for various ways to increase the complexity. Example 2 represents a small step up in complexity by replacing the fixed, class-specific *k*-mer with a class-specific position weight matrix (PWM), which is a common representation of naturally occurring sequence elements, such as binding sites for a transcription factors. Another small step up in complexity is example 3, where the PWM is placed randomly within in sequence window. In example 4 none of the PWMs is class-specific, only a combination of PWMs. Rules like these could be used to model cell type specific chromatin accessibility that is dependent on the interaction between transcription factors. Examples 5 and 6 encode class information in the relative position of PWMs instead of their presence or absence, with example data set 5 using class-specific order constraints and example data set 6 class-specific spacing constraints.

Once the synthetic data set is generated, it is used by the learner component in step 2 to train a neural network model. It is important to note that the learner only has access to *x* and *y* of the (*x, y, a*) example triplets, and the positional annotations *a* are only utilized in step 3. Analogous to the role of the data definition for the simulator in step 1, the model definition (see Figure 1C) serves as a blueprint for the learner by providing a precise description of the neural network architecture, the loss function, the optimizer, and hyperparameters of the training process, and thus ensuring a reproducible model creation, training, and serving process for both PyTorch and TensorFlow models.

In step 3, the fully trained model from step 2 is then evaluated with the help of an array of conventional test set metrics and feature importance evaluators, such as Integrated Gradients [14] and Sufficient Input Subsets [18].

As a means to illustrate the various inputs and output of this pipeline, we prepared the results of a single seqgra analysis in Supplementary Figure S2. For this example, we used a simple grammar, similar to the one described in example 1 of Figure 1D, but instead of always inserting the class-specific *k*-mer, we use different insertion probabilities for each class, ranging from 100 % present in examples of class 1, *C*_1_, to 80 % present in *C*_2_, 60 %present in *C*_3_, 40 % present in *C*_4_, 20 % present in *C*_5_, 10 % present in *C*_6_, 5 % present in *C*_7_, and only present in 1 % of *C*_8_ examples. We chose a neural network architecture with two hidden layers, a convolutional layer, followed by a fully connected layer (Supplementary Figure S2A). After the simulation process finished, diagnostic plots were generated, depicting a heatmap of grammar positions for all examples per class (Supplementary Figure S2B). These so-called positional grammar probabilities (i.e., the probability for a specific position to be a grammar position), depicted in the heatmap correspond to the insertion probabilities of the grammar, as expected. Furthermore, the class-specific ROC curves in Supplementary Figure S2C show that the chosen neural network architecture was optimal in terms of predictive performance, with true positive rates of 1.0, 0.8, 0.6, 0.4, 0.2, 0.1, 0.05, and 0.01 (at the zero false positive level) for the classes *C*_1_ to *C*_8_, which are the theoretical upper limits given the insertion probabilities of the underlying grammar. This is also reflected in the precision-recall curves in Supplementary Figure S2D. In panels E to G we show the results of the feature importance evaluators raw gradient, absolute gradient, and Sufficient Input Subsets (see sections II-I and II-J for details). These heatmaps show whether the model’s predictions were based on relevant (i.e., grammar) positions and are therefore an indication of the model’s ability to recover the underlying grammar of the data set. All three methods suggest high grammar recovery (many dark green positions, few dark red positions).

Supplementary Figure S2 covered the results obtained from a single seqgra call, evaluating one neural network model trained on one synthetic data set, but most seqgra analyses compare various different architectures across a range of data sets (of different grammar complexities and sizes). For these situations, we provide a suite of convenient commands that streamline these analyses and provide a schematic description of their inputs and outputs in Supplementary Figure S1.

### B. Selection of unambiguous set of HOMER sequence motifs

In order to generate synthetic data sets that are closer to experimentally obtained data sets, we replaced the artificially constructed *k*-mers used in the insertion probability grammar of Supplementary Figure S1 with transcription factor binding site motifs which were obtained from ChIP-seq assays and curated by HOMER [10]. However, before a collection of experimentally obtained motifs can be used effectively as sequence elements in grammars, degenerate motifs must be excluded. These include motifs with low information content and highly similar motif pairs. If these motifs are used as sequence elements that encode information about the condition *y*, but either cannot be differentiated from the background distribution or motifs specific to one condition are highly similar to motifs specific to another condition, the conditions are rendered inseparable and learning becomes impossible. This scenario is shown in Figure 2A, which depicts the test set ROC curves of a Bayes Optimal Classifier (BOC) for 10 classes of a data set generated by a grammar using 10 randomly selected HOMER motifs as class-specific sequence elements. BOCs in the context of seqgra are used to determine whether the conditions of a grammar are separable in principle, i.e., regardless of data set size and neural network architecture. Instead of neural network models whose weights are adjusted during training, the BOC has access to the data definition and uses the rules and sequence elements specified there directly to classify the examples. If the predictive performance of the BOC is low, as is the case with conditions *C*_6_, *C*_8_,and *C*_4_ shown in Figure 2A, the rules associated with those conditions are not specific enough to differentiate between them. And since the rules in this case place a supposedly condition-specific sequence element at a random position in the sequence window, the only explanation is that these sequence elements are either indistinguishable from background or indistinguishable from each other. The latter is shown in the matrices in Figure 2C and Figure 2D, which identify the corresponding sequence elements SE_6_, SE_8_, and SE_4_ as most similar to other sequence elements, i.e., lowest KL divergence and highest empirical similarity score, respectively (for details, see sections II-D and II-E).

**Fig. 2.**
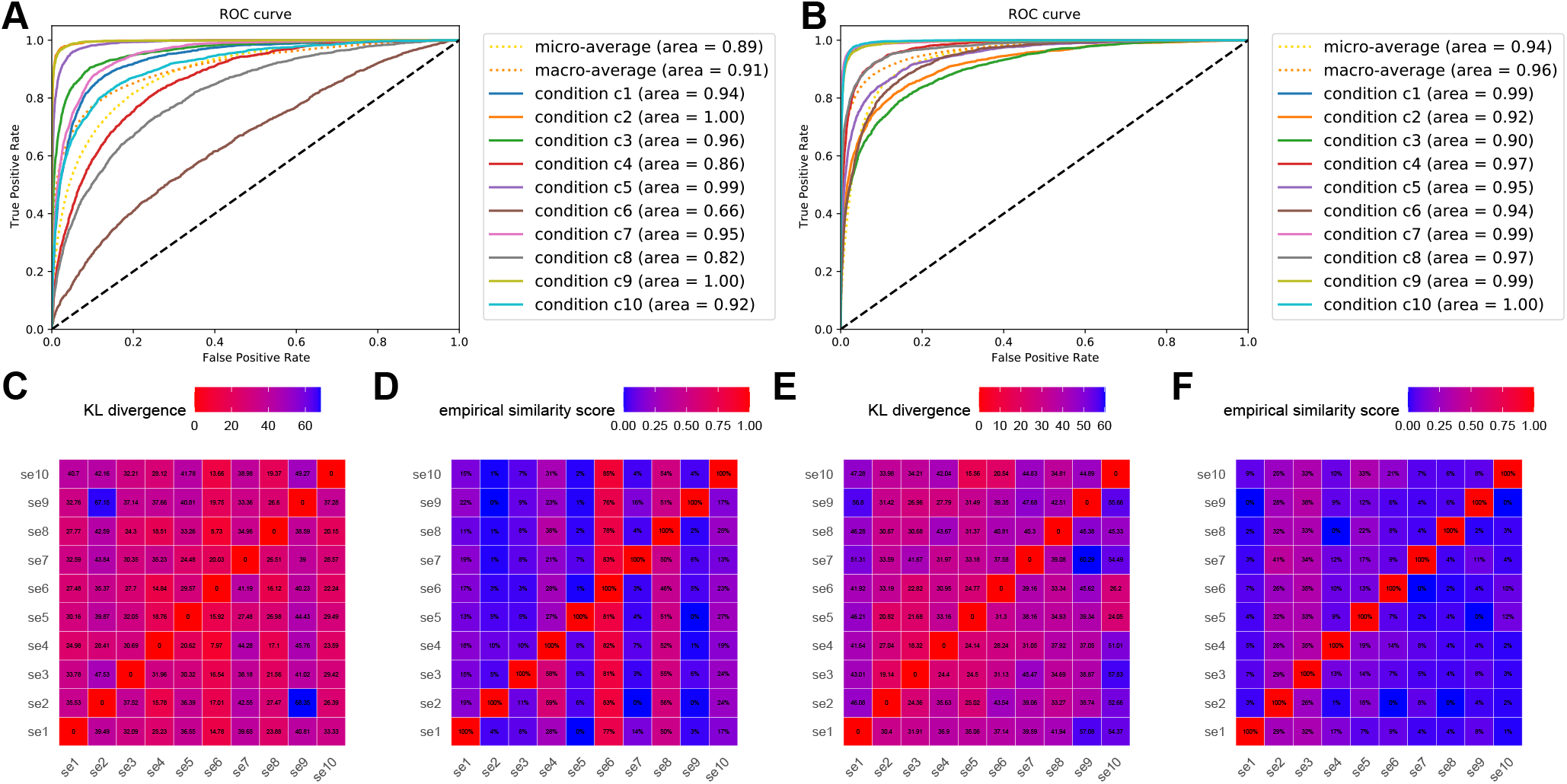
Selection of sequence motifs for simulation grammars. (**A**) ROC curve of Bayes Optimal Classifier on multi-class classification task with 10 classes, prior to filtering out ambiguous sequence motifs. (**B**) Same as panel A, after ambiguous sequence motifs were removed. (**C**) KL divergence matrix of 10 sequence motifs, prior to filtering. (**D**) Empirical similarity score matrix of 10 sequence motifs, prior to filtering. (**E**) Same as panel C, after removing ambiguous motifs. (**F**) Same as panel D, after removing ambiguous motifs.

Figure 2B shows BOC performance after the most ambiguous motifs were removed, and the corresponding KL divergence and empirical similarity score matrices are shown in Figure 2E and F. A collection of experimentally derived sequence motifs will never be completely orthogonal, but the degree of dissimilarity between these 10 were deemed sufficient and all subsequent multi-class classification grammars with 10 classes used these 10 motifs. Supplementary Figure S3 shows the same selection process for a collection of 100 HOMER motifs. All HOMER motifs used in this study are listed in Supplementary Table S1, together with a IUPAC notation of the motif, the motif information content (see section II-B) and the KL divergence between the motif and the background distribution (see section II-C). Motifs used for binary classification tasks are listed in Supplementary Table S2, those for multi-class classification tasks with 10, 20, and 50 classes are listed in Supplementary Tables S3, S4, and S5, respectively.

### C. seqgra-enabled ablation analysis reveals most efficient neural network architecture

Ablation, a technique widely used in neuroscience to determine the functions of brain regions by removing them one by one, has been used similarly to identify the relevant components of an artificial neural network [19,20]. We performed an ablation analysis to determine the effects of dropout [21] and batch normalization [22] on the predictive performance and grammar recovery of a basic neural network architecture with two hidden layers, a convolutional layer with 10 21-nt wide filters, followed by a dense layer with 5 hidden units, and dropout or batch normalization operations after each layer. Models were trained on binary classification data sets generated by grammars using class-specific HOMER motifs (see schematic in Figure 3A), class-specific order of HOMER motifs (Figure 3B), and class-specific spacing constraints between HOMER motifs (Figure 3C). Test set precision-recall curve AUCs are shown for all models across all grammars in Figure 3D. Unsurprisingly, the predictive performance of all architectures increases with data set size, and all architectures approach a PR AUC of 1.0 for sufficiently large data sets. But this analysis reveals a striking difference between the neural network architectures in terms of their efficiency, i.e., how many training examples are required to reach an AUC of approximately 1.0. On the grammars tested here, batch normalization had a negative effect on efficiency, requiring up to 100,000 examples more to converge than architectures without the operation. The architecture with dropout after each hidden layer was the most efficient and highest performing, both in terms of predictive performance and grammar recovery (i.e., the model’s propensity to classify examples based on grammar positions) as shown in Figure 3E.

**Fig. 3.**
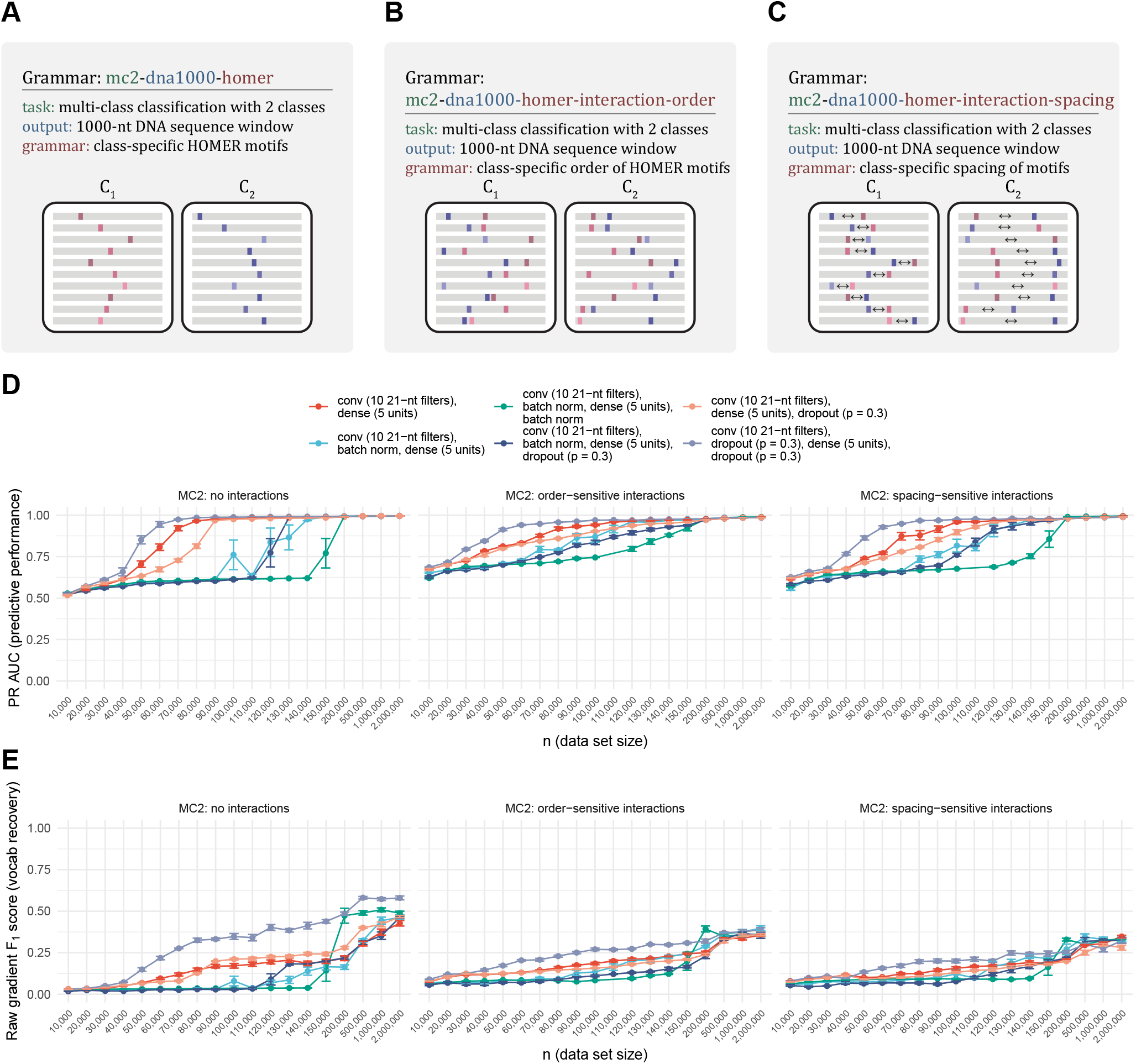
seqgra-enabled ablation analysis reveals most efficient neural network architecture. (**A**) Schematic of binary classification grammar using class-specific HOMER motifs as sequence elements. (**B**) Schematic of binary classification grammar using class-specific order of HOMER motifs. (**C**) Schematic of binary classification grammar using class-specific spacing of HOMER motifs. (**D**) Predictive performance of six neural network architectures with and without batch normalization and dropout. (**E**) Vocabulary recovery of six neural network architectures with and without batch normalization and dropout.

### D. DeepSEA dominates comparison of popular genomics deep learning architectures

Furthermore, we compared three popular neural network architectures used in the field of genomics, Basset [3], ChromDragoNN [6], and DeepSEA [5]. All three architectures were devised with functional genomics data sets in mind and were originally trained on multi-label classification data sets obtained from numerous DNase-seq assays, with ChromDragoNN also utilizing RNA-seq and DeepSEA ChIP-seq data. With over 4 million (Basset), over 6 million (DeepSEA), and over 20 million (ChromDragoNN) trainable parameters, all three can be considered high-capacity models. The three architectures make use of commonly used building blocks such as convolutional, followed by dense layers (all three), max pooling and dropout operations (all three), ReLU activation functions (all three), batch normalization (Basset and ChromDragoNN), and skip connections (ChromDragoNN). Input and output layers were adjusted to fit the prediction task and architectures were trained on simulated data sets from scratch without pre-training on their original data sets.

We used the area under the micro-averaged precision-recall curve to evaluate the test set predictive performance on four multi-class classification tasks (with 2, 10, 20, and 50 classes) and three or four grammars each, with a sequence window of 1000 nucleotides. The results are shown in Figure 4A for binary classification, and Figures 4B, 4C, and 4D for multi-class classification with 10, 20, and 50 classes, respectively. The HOMER motifs used by the grammars presented here are listed in Supplementary Tables S2-S5. Each panel contains precision-recall AUCs of models trained on data sets generated by one grammar, using 5 different random seeds for simulation (error bars) and 19 different data set sizes. The DeepSEA architecture exhibited an at times substantially higher predictive performance than Basset and ChromDragoNN and was the highest performing architecture on all tested data sets. While DeepSEA is the preferred architecture on data sets derived from the grammars we tested, this is not necessarily true for data sets with other grammars or experimentally obtained data. Interestingly, we observed that high capacity architectures such as those tested here perform better on data sets generated by grammars that include interactions, specifically interactions that encode the class label in the order or spacing of the interacting sequence elements. This is not the case for small-scale architectures with less than 100,000 trainable parameters, which, as expected, do better on grammars without interactions, where the class label is encoded in the presence of class-specific sequence elements.

**Fig. 4.**
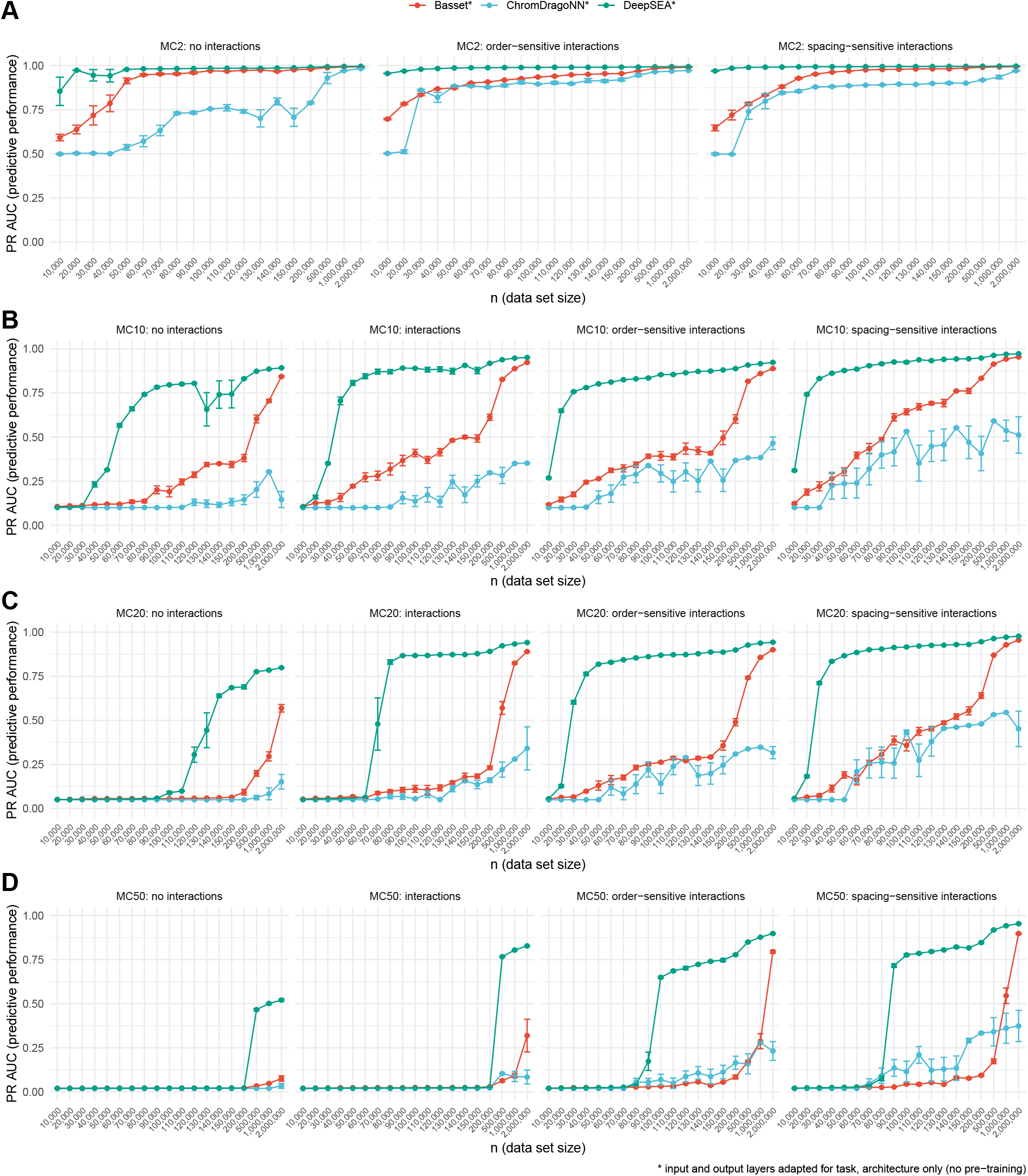
Comparison of neural network architectures Basset, ChromDragoNN, and DeepSEA. (**A**) Predictive performance on binary classification tasks of grammars with class-specific HOMER motifs (left), class-specific order of HOMER motifs (middle), and class-specific spacing of HOMER motifs. All architectures were trained on data sets ranging in size from 10,000 examples to 2,000,000 examples. Error bars are standard errors of five models trained on the same grammar, using five different simulation seeds. (**B**) Same as panel A, for multi-class classification tasks with 10 classes. The second plot from the left shows the predictive performance of models trained on data sets with class-specific interactions of HOMER motifs. (**C**) Same as panel B, for multi-class classification tasks with 20 classes. (**D**) Same as panel B, for multi-class classification tasks with 50 classes.

### E. High predictive performance of simulation-vetted neural network architecture recapitulated with ChIP-seq data

In this section we address the question of whether neural network architectures that perform well on simulated data also succeed on data obtained experimentally. We decided to model the well-known hetero-dimeric pair of transcription factors SOX2 and POU5F1, whose spacing constraints were previously characterized [23,24]. To that end, we used the HOMER motifs `SOX2_HUMAN.H11MO.0.A` and `PO5F1_HUMAN.H11MO.1.A` as sequence elements in the data definition. We also included spacing constraints (0-3 bp between SOX2 and PO5F1 motifs). Figure 5A shows a schematic depiction of the analysis.

**Fig. 5.**
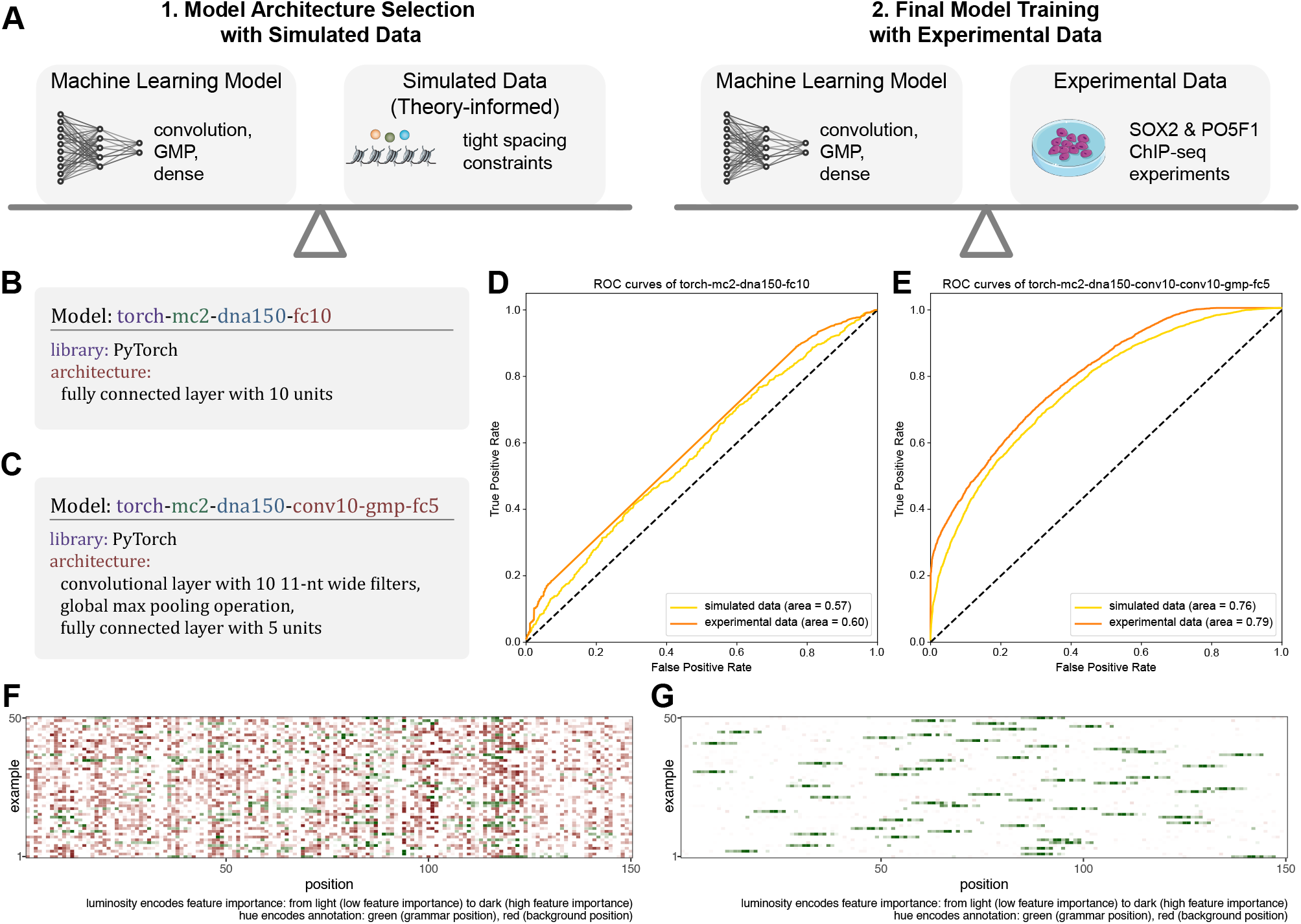
Predictive performance and grammar recovery of various model architectures on simulated and experimental data. (**A**) Schematic of model selection process: first, identify suitable model architectures on simulated data; second, train models with simulation-vetted architectures on experimental data. (**B**) Naive neural network architecture with fully connected layer. (**C**) Grammar-informed neural network architecture with convolutional layer, global max pooling, and fully connected layer. (**D**) Predictive performance of naive architecture, trained and evaluated on simulated and experimental data. (**E**) Predictive performance of grammar-informed architecture, trained and evaluated on simulated and experimental data. (**F**) Grammar agreement plot (Integrated Gradients) of naive architecture, trained on experimental data. (**G**) Grammar agreement plot (Integrated Gradients) of grammar-informed architecture, trained on experimental data.

The experimental data set was based on two ChIP-seq assays, which targeted the two transcription factors. The preprocessed data was obtained from the Cistrome Data Browser [25], specifically the data associated with GEO IDs `GSM1701825` for SOX2 and `GSM1705258` for POU5F1.

We evaluated the same neural network architectures on both the simulated and the experimental data sets. The architecture described in Figure 5B with one fully connected layer (not counting the output layer) is an example of an architecture that does not assume any structure in the input. It is a naive architecture in the sense that it was constructed without any knowledge about the grammar that was used to simulate the data. The architecture described in Figure 5C, on the contrary, makes assumptions about the data that are in agreement with the grammar, such as a 1D spatial structure with information encoded in 11-nt long code words (enough to cover the SOX2-POU5F1 interaction), whose position in the sequence window is irrelevant.

As expected, the test set predictive performance of the naive architecture (Figure 5D) was significantly lower than the grammar-informed architecture (Figure 5E). Furthermore, the performance on the simulated data proved to be a good predictor for the performance on the experimental data (Figure 5D and E).

The agreement between feature importance and the grammar positions, a proxy for a model’s ability to recover the SOX2 and POU5F1 motifs, is shown in Figure 5F for the naive architecture and in Figure 5G for the grammar-informed architecture. The grammar-informed model’s predictions were based almost exclusively on grammar positions (positions that contained SOX2 and POU5F1 motifs), whereas this was not the case for the naive model. Both panels were created with the Integrated Gradients feature importance evaluator.

## IV. Discussion

In this paper we introduced seqgra, a deep learning infrastructure method for genomics. It is intended to streamline the development of deep learning models for biological sequence-based prediction tasks, by providing a reproducible unified framework for (1) flexible, rule-based synthetic data generation; (2) model training; and (3) model evaluation with conventional test set metrics and feature attribution methods. This three-step pipeline supports data sets obtained by simulation and experiment, models implemented in PyTorch and TensorFlow, and numerous gradient-based feature attribution methods as well as Sufficient Input Subsets, a model-agnostic feature attribution method, in addition to conventional ROC and precision-recall curves for model evaluation. Our method greatly simplifies an array of commonly performed diagnostics and performance assessments of deep learning models, such as ablation analysis, estimated data set size requirements, and tolerated noise thresholds. The simulator and the language of the probabilistic rules are flexible enough to span multi-class and multi-label classification tasks with any number of classes or labels, DNA or amino acid sequence windows of variable or fixed length, class-dependent background distributions, sequence elements defined as position weight matrices or list of *k*-mers with associated probabilities, and interactions between sequence elements with associated order or spacing constraints.

Moreover, the controlled environment of data simulation and reproducible model training, serving, and evaluation makes seqgra a suitable testbed for feature attribution and interpretability methods and their interdependencies with neural network architectures and the complexity level of the training data. The framework can even be used to perform extensive comparisons between deep learning libraries, which are rarely done (see Supplementary Figures S7 and S8) or identify undocumented behavior of the deep learning technology stack, such as an unusual training instability caused by a random seed of zero on some grammar-architecture combinations, which is reproducible and occurs in both PyTorch and TensorFlow (see Supplementary Figures S4 - S6).

To avoid confusion, we would like to point out that seqgra is not a neural architecture search technique in the sense that it will not propose suitable neural network architectures for a particular data set. The model definition is an input, not an output of the seqgra pipeline. However, seqgra can be used in conjunction with neural architecture search, such as AMBER [26], a neural architecture search method for architectures aimed at genomics prediction tasks, or general hyperparameter optimization methods, such as Hyperband [27].

One caveat of all simulation-based approaches is the inevitable gap between simulated and real-world data sets, in the sense that the former is always a simplified approximation of the latter. Thus insights gained from simulated data might not carry over to the experimental world. In fact, to a certain degree this will always be the case. However, while high-performing neural network architectures on simulated data might not perform as highly on experimental data, the opposite is rarely the case, i.e., low-performing architectures in simulation are unlikely to improve when trained on noisier and/or smaller experimental data sets.

While the intricacies of noisy and biased high-throughput genomics experiments make for highly complex and poorly understood data sets, training highly complex alchemy-like [28] deep neural networks on them contributes little to a mechanistic understanding of the biological processes that are at work underneath and might worsen the reproducibility crisis in both machine learning [29] and biology [30,31]. Simulated data, however, is perfectly understood, its noise levels controlled and any biases artificially introduced and accounted for, which makes it an excellent environment for model evaluation. With seqgra, the clean room of simulated data and a precise description of the patterns in the data (i.e., the probabilistic rules in the data definition) on the one end is paired with an array of feature attribution methods on the other, to answer questions that are often impossible to answer with poorly understood genomics data. One such question is whether the predictions of the model are based on those parts of the input that are in fact relevant for the phenomenon that is predicted, or, to put it another way, whether the model was able to recover the underlying rules of the data set.

## Availability

The source code of the seqgra package is hosted on GitHub (https://github.com/gifford-lab/seqgra) and licensed under the MIT license. seqgra is part of the Python Package Index PyPI and can be installed using pip, the Python package installer. Extensive documentation can be found at https://kkrismer.github.io/seqgra.

## Author contributions

Conceptualization, K.K, D.K.G.; Methodology, K.K.; Software, K.K. and J.H; Formal Analysis, K.K. and J.H.; Investigation, K.K.; Resources, D.K.G.; Data Curation, K.K.; Writing - Original Draft, K.K.; Writing - Review & Editing, K.K., J.H., and D.K.G.; Visualization, K.K.; Supervision, D.K.G.; Funding Acquisition, D.K.G.

## Acknowledgements

We thank members of the Gifford lab for insightful suggestions and discussions.

