## Supplementary Material for "seqgra: Principled Selection of Neural Network Architectures for Genomics Prediction Tasks"

### 1 Supplementary Figures

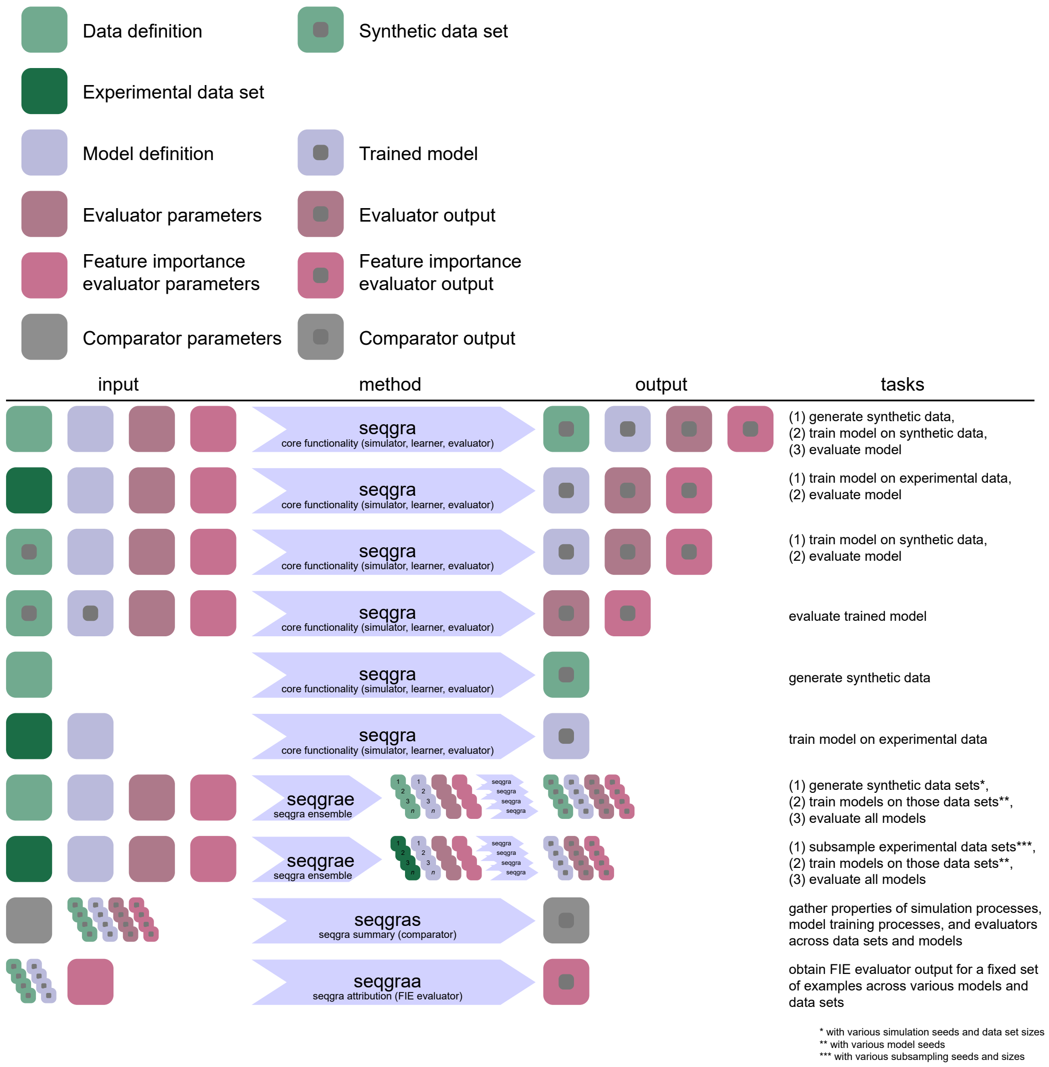


**Supplementary Figure S1: Schematic of common use cases for seqgra command line interface.** The seqgra package contains four commands, `seqgra`, `seqgrae`, `seqgras`, and `seqgraa`. `seqgra` contains the core functionality of (1) generating synthetic data using the *Simulator* component, (2) training models on either synthetic or experimental data using various *Learner* components, and (3) evaluating the model using various *Evaluator* components. `seqgrae`, short for seqgra ensemble, is a convenient way to generate multiple synthetic data sets with various data set sizes and simulation seeds and train models on them using a range of model seeds. `seqgras`, short for seqgra summary, is a tool to gather properties and metrics across a number of data sets and models and compare them using *Comparator* components. `seqgraa`, short for seqgra attribution, runs feature importance evaluators on a number of trained models, using the same set of examples each time.

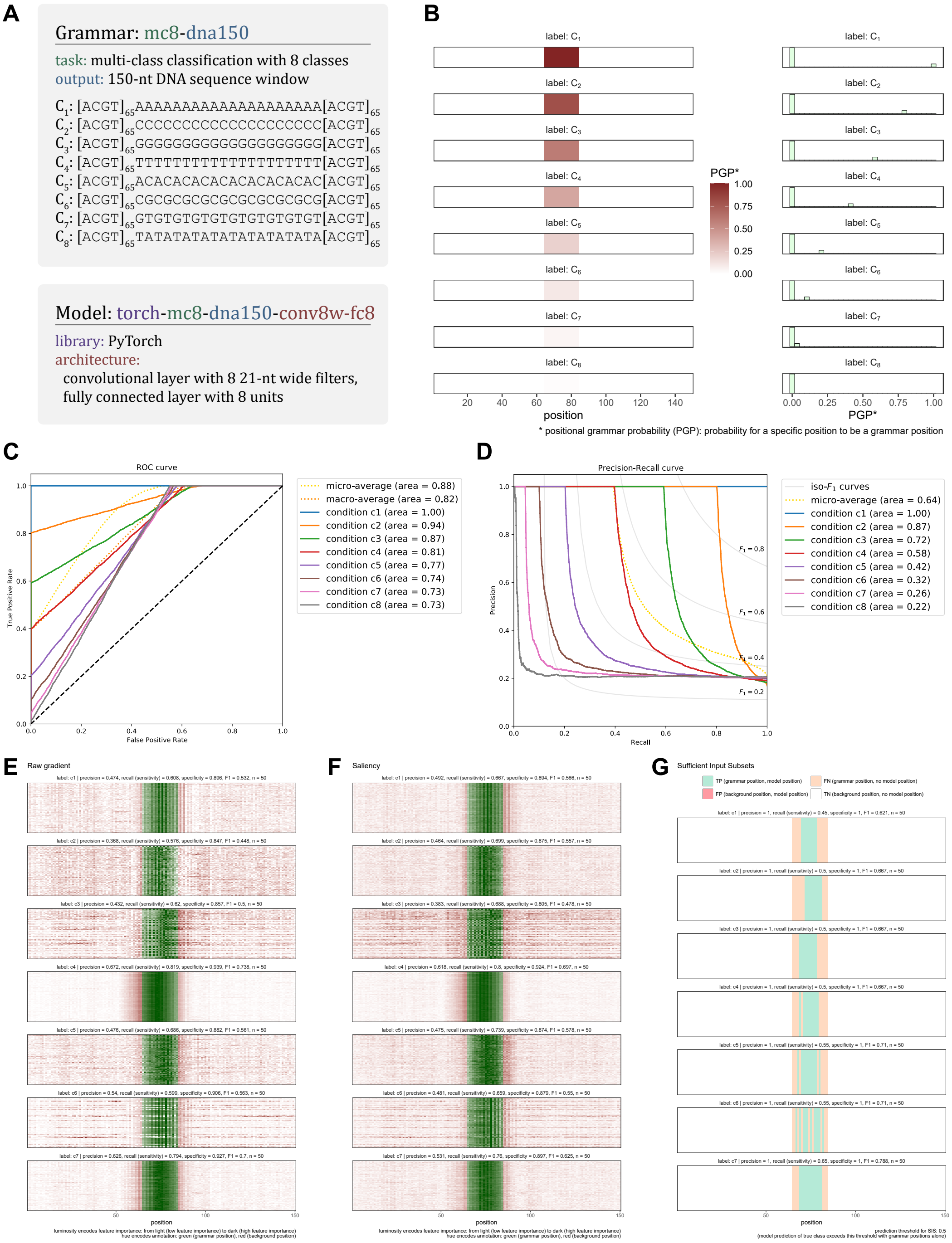


**Supplementary Figure S2: Insertion probability test.** (**A**) Grammar and model description. (**B**) Grammar position heatmap (on the left) depicting the probability of grammar annotation for all positions (1 - 150) and all classes ($C_1$ to $C_8$). (**C**) Test set ROC curve of classifier trained on synthetic data depicts class-specific true positive rates that mirror insertion probabilities, as expected. (**D**) Test set PR curve of same classifier, class-specific curves mirror insertion probabilities. (**E**) Raw gradient feature importance for classes $C_1$ to $C_7$ (classifier did not correctly predict class $C_8$). The x-axis is the position in the sequence window, the y-axis are randomly drawn examples with that class label. Dark green areas are grammar positions with high feature importance (desired), dark red areas are background positions with high feature importance (undesired). (**F**) Same as panel E, for absolute gradient (saliency) feature importance. (**G**) Same as panel E, for Sufficient Input Subsets (SIS) feature importance. The only difference is that SIS is an inherently discrete measure of feature importance, either positions are part of a sufficient input subset or not.

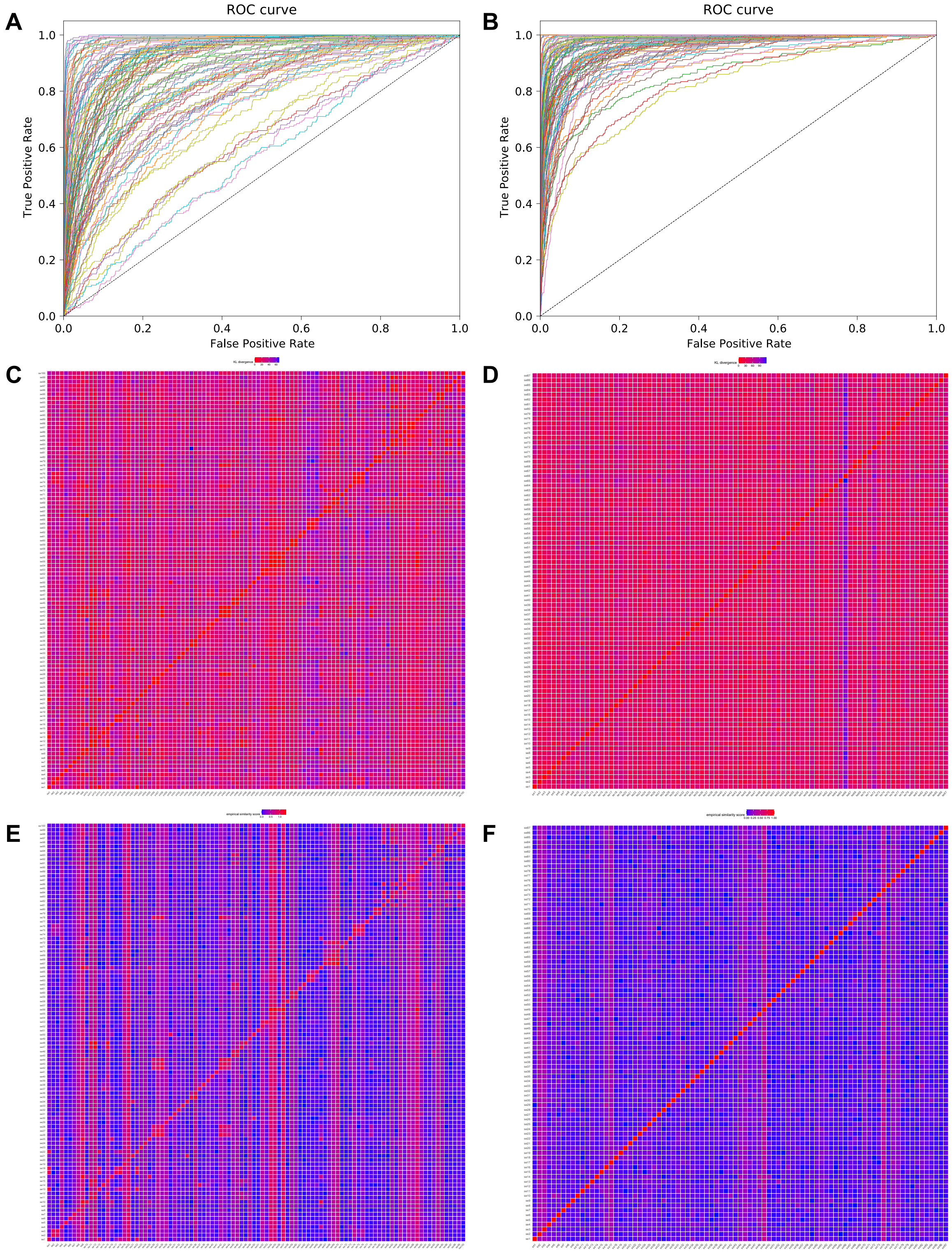


**Supplementary Figure S3: Selection of sequence motifs for MC100 simulation grammars.** (**A**) ROC curve of Bayes Optimal Classifier on multi-class classification task with 100 classes, prior to filtering out ambiguous sequence motifs. (**B**) Same as panel A, after ambiguous sequence motifs were removed. (**C**) KL divergence matrix of 100 sequence motifs, prior to filtering. (**D**) Same as panel C, after removing ambiguous motifs. (**E**) Empirical similarity score matrix of 100 sequence motifs, prior to filtering. (**F**) Same as panel D, after removing ambiguous motifs.

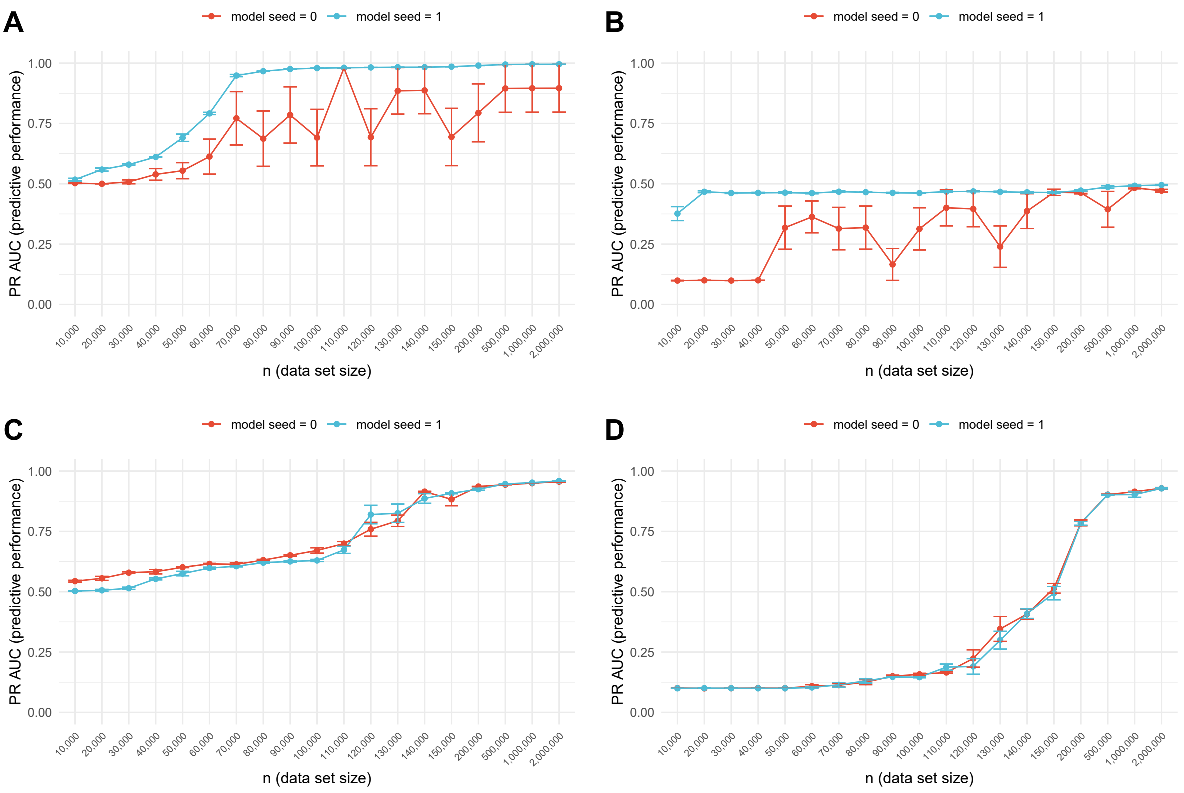


**Supplementary Figure S4: PyTorch and TensorFlow affected by random seed induced instability.** (**A**) Shown are test set PR AUCs of a PyTorch neural network architecture with two hidden layers, a convolutional layer with 10 21-nt wide filters followed by a dense layer with 5 units, trained on binary classification data sets using HOMER motifs without interactions. This PyTorch neural network architecture exhibits an unusual variability in PR AUC when trained with a random seed of zero. (**B**) Shown are test set PR AUCs of a TensorFlow neural network architecture with a convolutional layer with 10 21-nt wide filters, a global max pooling operation, and a dense layer with 10 units, trained on multi-class classification data sets with 10 classes using HOMER motifs with spacing-sensitive interactions. This TensorFlow neural network architecture exhibits an unusual variability in PR AUC when trained with a random seed of zero. (**C**) Unlike panel A, this PyTorch neural network architecture (convolutional layer with 10 11-nt wide filters followed by a dense layer with 5 units), which was trained on the same data sets as the one in panel A, does not exhibit unusually high variability of PR AUCs when trained with a random seed of zero. (**D**) Similarly, this TensorFlow neural network architecture (convolutional layer with 10 21-nt wide filters followed by a dense layer with 10 units), which was trained on the same data sets as the one in panel B, also does not exhibit unusually high variability of PR AUCs when trained with a random seed of zero.

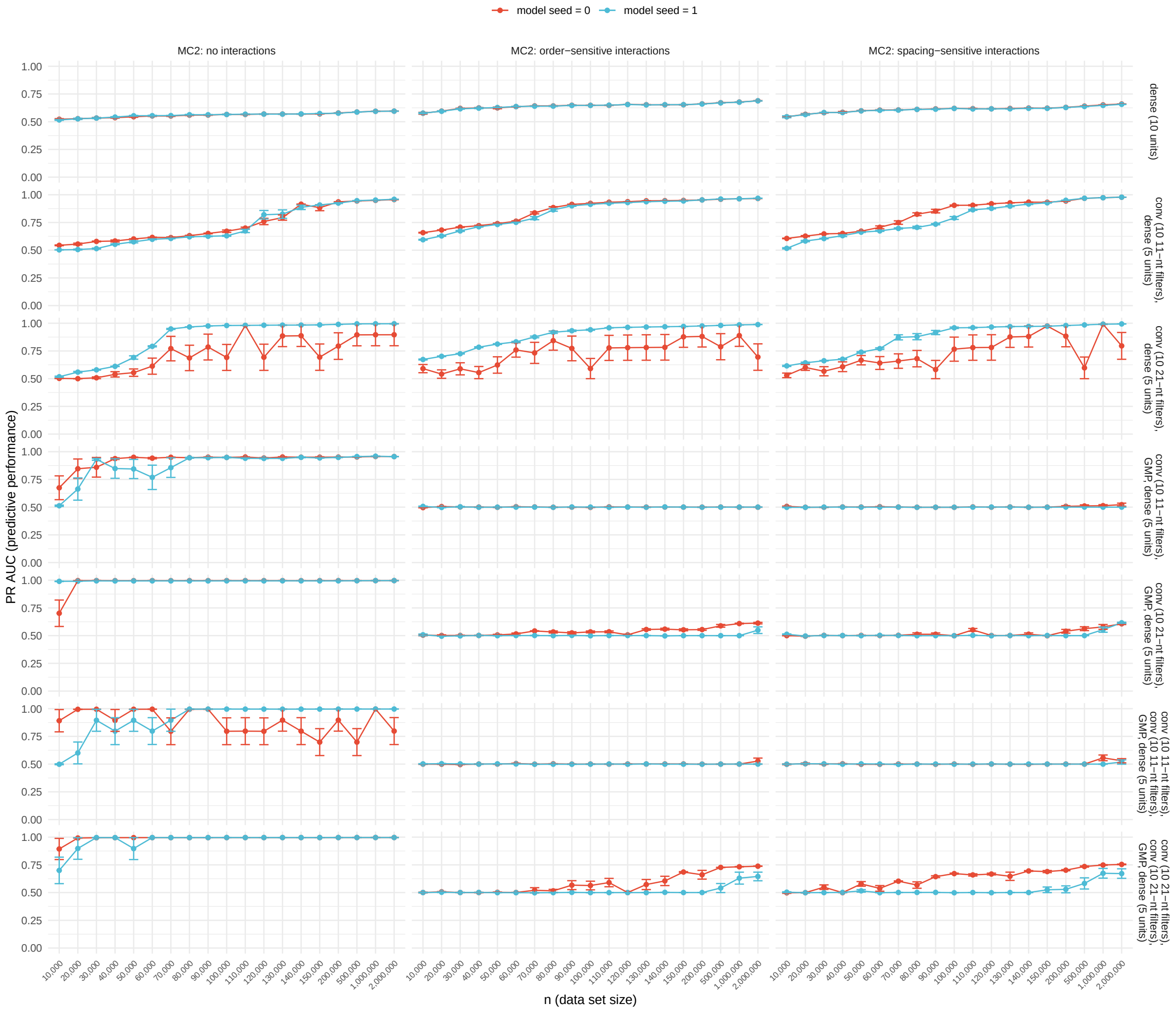


**Supplementary Figure S5: PyTorch models trained with random seed 0 suffer from grammar-dependent and architecture-dependent instability.** Not all grammar-architecture combinations are affected.

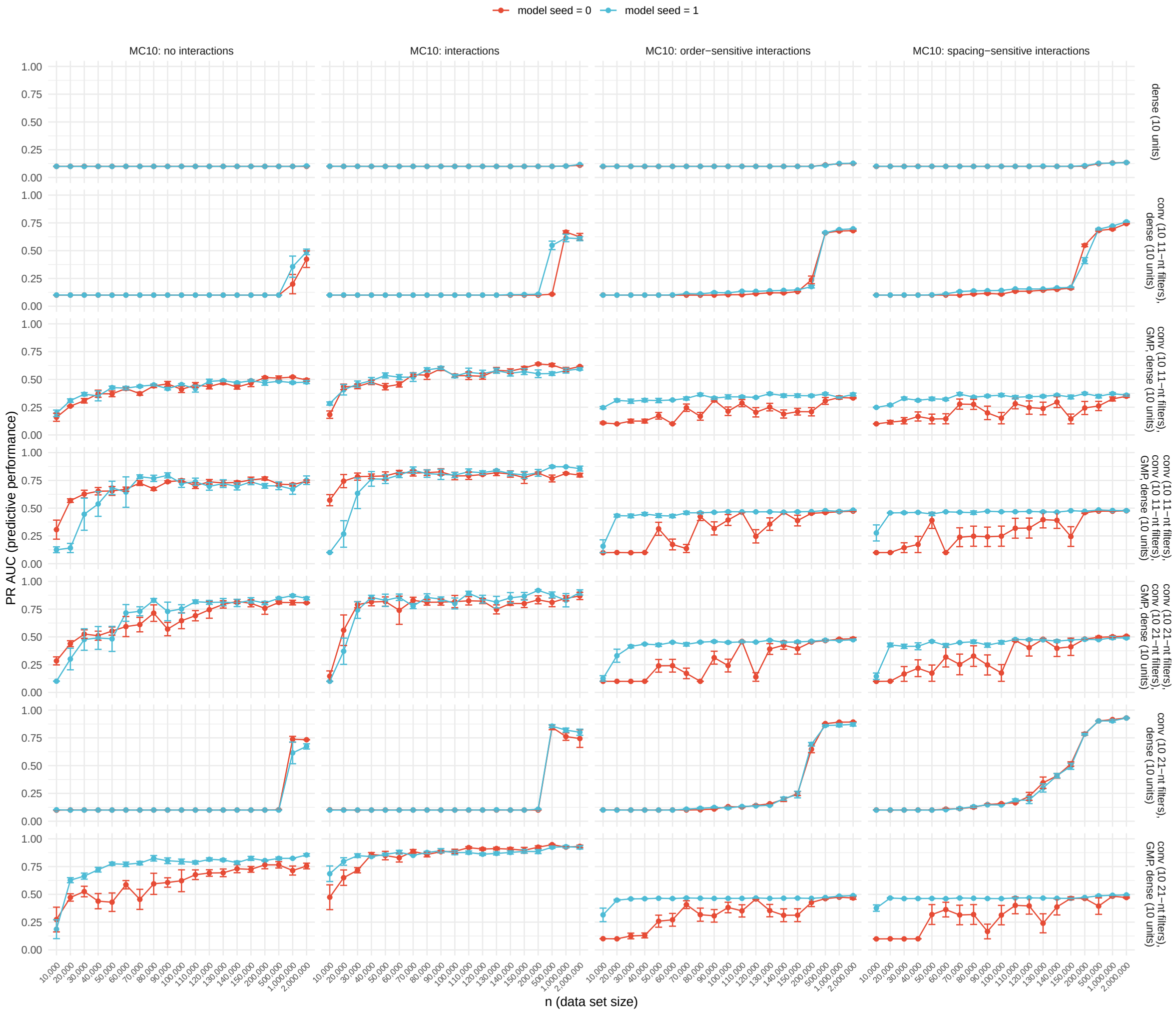


**Supplementary Figure S6: TensorFlow models trained with random seed 0 suffer from grammar-dependent and architecture-dependent instability.** Not all grammar-architecture combinations are affected.

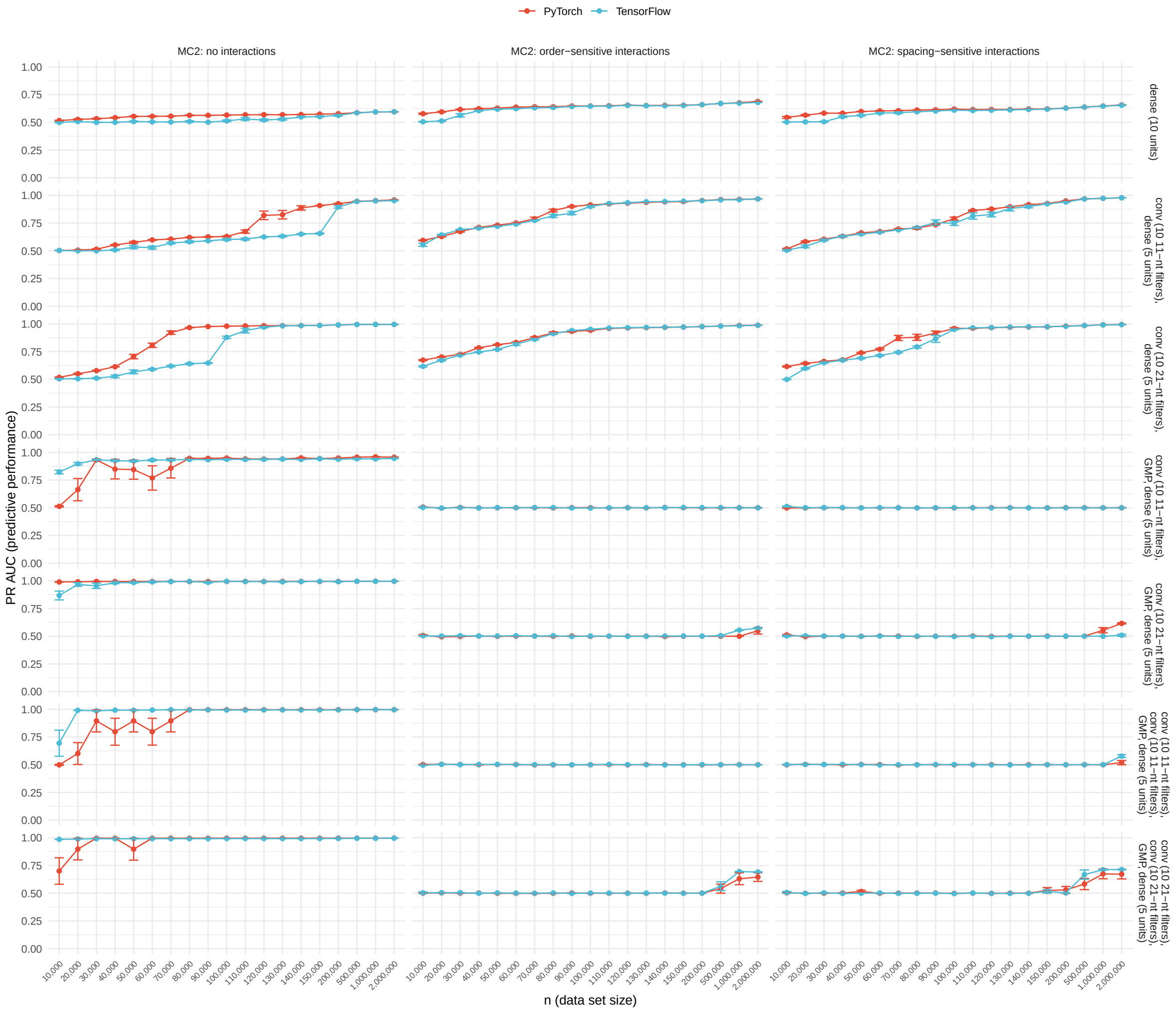


**Supplementary Figure S7: Comparison of PyTorch and TensorFlow architectures trained on binary classification data sets.** When comparing an equivalent neural network architecture, trained on the same data set, test set PR AUCs between models implemented and trained with deep learning libraries PyTorch and TensorFlow are similar. Shown here are comparisons across three grammars, 19 data set sizes, and seven neural network architectures.

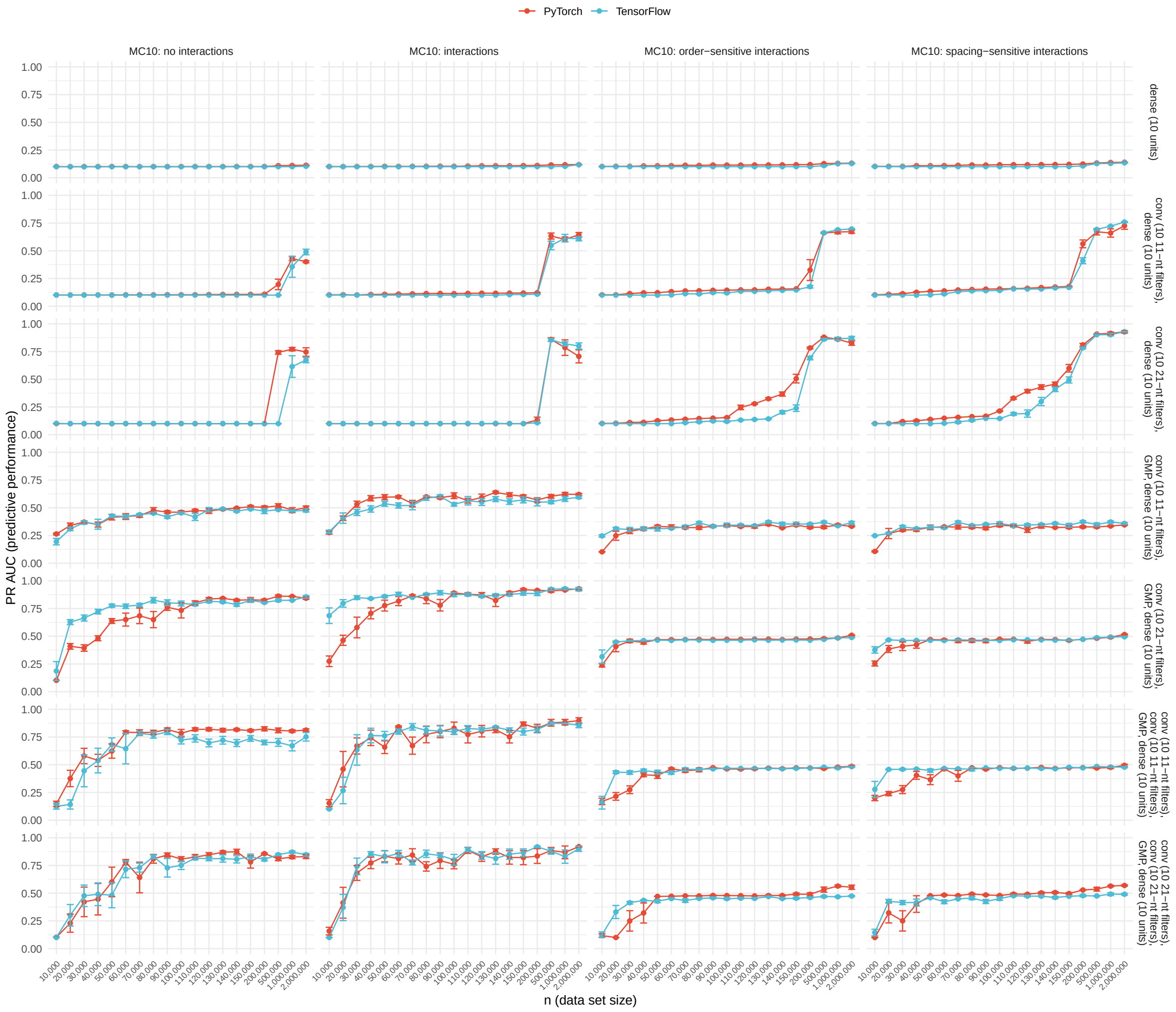


**Supplementary Figure S8: Comparison of PyTorch and TensorFlow architectures trained on multi-class classification data sets with 10 classes.** When comparing an equivalent neural network architecture, trained on the same data set, test set PR AUCs between models implemented and trained with deep learning libraries PyTorch and TensorFlow are similar. Shown here are comparisons across four grammars, 19 data set sizes, and seven neural network architectures.

#### 2 Supplementary Tables

| ID | Motif | IUPAC notation | Width (in nt) | MIC | $D_{KL}(\cdot)$ |
|---|---|---|---|---|---|
| se1 | FOXA1:AR | AGTAAACAAAAAAGAACANA | 20 | 18.9 | 17.4 |
| se2 | Bcl11a | TYTGACCASWRG | 12 | 11.3 | 11.8 |
| | | MC2 grammar motifs above | | | |
| se3 | Brachyury | ANTTMRCASBNNNGTGYKAAN | 21 | 11.5 | 11.6 |
| se4 | CEBP:CEBP | NTNATGCAAYMNNHTGMAAY | 20 | 14.8 | 14.3 |
| se5 | Chop | ATTGCATCAT | 10 | 13.2 | 12.7 |
| se6 | CHR | CGGTTTCAAA | 10 | 12.6 | 11.8 |
| se7 | CTCF-SatelliteElement | TGCAGTTCCAANAGTGGCCA | 20 | 18.8 | 19.6 |
| se8 | Mouse Recombination Hotspot | ACTYKNATTCGTGNTACTTC | 20 | 15.3 | 14.9 |
| se9 | RAR:RXR | RGGTCADNNAGAGGTCAV | 18 | 16.3 | 17.3 |
| se10 | DUX | BCWGATTCAATCAAN | 15 | 17.9 | 16.9 |
| | | MC10 grammar motifs above | | | |
| se11 | E2F7 | VDTTTCCCGCCA | 12 | 13.4 | 14.6 |
| se12 | EBNA1 | GGYAGCAYDTGCTDCCCNNN | 20 | 18.1 | 19.2 |
| se13 | ERE | AAGGTCACNGTGACC | 15 | 14.3 | 15.2 |
| se14 | ETS:E-box | AGGAAACAGCTG | 12 | 17.3 | 17.6 |
| se15 | EWS:ERG-fusion | ATTTCCTGTN | 10 | 13.7 | 13.5 |
| se16 | Foxh1 | NNTGTGGATTSS | 12 | 11.3 | 11.1 |
| se17 | FXR | AGGTCANTGACCTN | 14 | 12.3 | 13.2 |
| se18 | GATA3 | AGATGKDGAGATAAG | 15 | 17.3 | 16.5 |
| se19 | GATA3 | AGATSTNDNNDSAGATAASN | 20 | 16.9 | 16.3 |
| se20 | GATA | NAGATWNBNATCTNN | 15 | 14.0 | 13.3 |
| | | MC20 grammar motifs above | | | |
| se21 | GATA:SCL | CGGCTGCNGNNNNCAGATAA | 20 | 15.4 | 15.9 |
| se22 | Gfi1b | AAATCACTGC | 10 | 13.9 | 13.8 |
| se23 | GRHL2 | AAACYKGTTWDACMRGTTTB | 20 | 13.5 | 13.4 |
| se24 | Hand2 | TGACANARRCCAGRC | 15 | 13.2 | 13.6 |
| se25 | HINFP | TWVGGTCCGC | 10 | 11.7 | 13.2 |
| se26 | HOXB13 | TTTTATKRGG | 10 | 13.5 | 12.6 |
| se27 | LRF | AAGACCCYYN | 10 | 11.2 | 12.5 |
| se28 | LXRE | GGGTTACTANAGGTCA | 16 | 17.5 | 17.9 |
| se29 | NF1:FOXA1 | NNTGTTTATTTTGGCA | 16 | 17.3 | 16.7 |
| se30 | NFAT:AP1 | GAATGGAAAAAATGAGTCAT | 20 | 15.5 | 15.1 |
| se31 | NFAT | ATTTTCCATT | 10 | 13.1 | 12.5 |
| se32 | NFY | AGCCAATCGG | 10 | 13.3 | 13.8 |
| se33 | Nur77 | TGACCTTTNCNT | 12 | 15.1 | 14.8 |
| se34 | Oct2 | ATATGCAAAT | 10 | 15.3 | 14.1 |
| se35 | Oct4:Sox17 | CCATTGTATGCAAAT | 15 | 15.9 | 15.0 |
| se36 | OCT4-SOX2-TCF-NANOG | ATTTGCATAACAATG | 15 | 16.4 | 14.9 |
| se37 | p53 | ACATGCCCGGGCAT | 14 | 16.7 | 18.2 |
| se38 | PAX3:FKHR-fusion | ACCGTGACTAATTNN | 15 | 14.6 | 14.1 |

*Continued on next page*

Supplementary Table S1 – *Continued from previous page*

| ID | Motif | IUPAC notation | Width (in nt) | MIC | $D_{\mathrm{KL}}(\cdot)$ |
|---|---|---|---|---|---|
| se39 | PAX5 | `GTCACGCTCNCTGA` | 14 | 15.1 | 16.3 |
| se40 | PAX6 | `NGTGTTCAVTSAAGCGKAAA` | 20 | 13.9 | 14.3 |
| se41 | Pax7 | `NTAATTDGCYAATTANNWWD` | 20 | 16.0 | 13.9 |
| se42 | Pax7 | `TAATCAATTA` | 10 | 16.3 | 14.6 |
| se43 | Pax8 | `GTCATGCHTGRCTGS` | 15 | 13.4 | 14.6 |
| se44 | Pitx1:Ebox | `YTAATTRAWWCCAGATGT` | 18 | 12.7 | 11.8 |
| se45 | PRDM10 | `TGGTACATTCCA` | 12 | 11.9 | 12.3 |
| se46 | PRDM14 | `AGGTCTCTAACC` | 12 | 13.7 | 14.0 |
| se47 | PRDM15 | `YCCDNTCCAGGTTTT` | 15 | 13.2 | 13.7 |
| se48 | PRDM9 | `ADGGYAGYAGCATCT` | 15 | 12.8 | 13.1 |
| se49 | PSE | `WAVTCACCMTAASYDAAAAG` | 20 | 10.6 | 10.3 |
| se50 | RBPJ:Ebox | `GGGRAARRGRMCAGMTG` | 17 | 14.3 | 15.2 |

**Supplementary Table S1: Homer transcription factor motifs for multi-class classification tasks with 2, 10, 20, and 50 classes (MC2-MC50):** These motifs are used for grammars without interactions, with interactions, with interactions with order constraints, and with interactions with spacing constraints. The columns (from left to right) contain the seqgra-internal sequence element ID, the motif name (name of the transcription factor or complex), a summary of the motif in IUPAC notation, the width of the motif in nucleotides, the motif information content (MIC), and the KL divergence between the motif and the background sequence (using the human genomic nucleotide distribution).

| ID | Motif | min $D_{\mathrm{KL}}(\cdot,\cdot)$ | max ESS$(\cdot,\cdot)$ |
|---|---|---|---|
| se1 | FOXA1:AR | 46.1 | 0 % |
| se2 | Bcl11a | 30.4 | 33 % |

**Supplementary Table S2: Homer transcription factor motifs for binary classification tasks (MC2):** These motifs are used for MC2 grammars without interactions, with interactions, with interactions with order constraints, and with interactions with spacing constraints. The columns (from left to right) contain the seqgra-internal sequence element ID, the motif name (name of the transcription factor or complex), the minimum KL divergence between the motif and the other motif in this grammar, and the maximum adjusted empirical similarity score (ESS) between the motif and the other motif in this grammar.

| ID | Motif | min $D_{\mathrm{KL}}(\cdot,\cdot)$ | max ESS$(\cdot,\cdot)$ |
|---|---|---|---|
| se1 | FOXA1:AR | 41.6 | 9 % |
| se2 | Bcl11a | 19.1 | 41 % |
| se3 | Brachyury | 18.3 | 38 % |
| se4 | CEBP:CEBP | 24.4 | 17 % |
| se5 | Chop | 15.6 | 33 % |
| se6 | CHR | 20.5 | 21 % |
| se7 | CTCF-SatelliteElement | 31.1 | 8 % |
| se8 | Mouse Recombination Hotspot | 33.3 | 6 % |
| se9 | RAR:RXR | 37.0 | 11 % |
| se10 | DUX | 24.1 | 12 % |

**Supplementary Table S3: Homer transcription factor motifs for multi-class classification tasks with 10 classes (MC10):** These motifs are used for MC10 grammars without interactions, with interactions, with interactions with order constraints, and with interactions with spacing constraints. The columns (from left to right) contain the seqgra-internal sequence element ID, the motif name (name of the transcription factor or complex), the minimum KL divergence between the motif and the other 9 motifs in this grammar, and the maximum adjusted empirical similarity score (ESS) between the motif and the other 9 motif in this grammar.

| ID | Motif | min $D_{\mathrm{KL}}(\cdot,\cdot)$ | max ESS$(\cdot,\cdot)$ |
|---|---|---|---|
| se1 | FOXA1:AR | 39.0 | 10 % |
| se2 | Bcl11a | 18.0 | 40 % |
| se3 | Brachyury | 18.3 | 35 % |
| se4 | CEBP:CEBP | 24.4 | 13 % |
| se5 | Chop | 15.6 | 32 % |
| se6 | CHR | 20.5 | 19 % |
| se7 | CTCF-SatelliteElement | 31.1 | 10 % |
| se8 | Mouse Recombination Hotspot | 24.3 | 9 % |
| se9 | RAR:RXR | 35.1 | 9 % |
| se10 | DUX | 24.1 | 14 % |
| se11 | E2F7 | 20.1 | 15 % |
| se12 | EBNA1 | 35.0 | 13 % |
| se13 | ERE | 16.8 | 18 % |
| se14 | ETS:E-box | 33.1 | 17 % |
| se15 | EWS:ERG-fusion | 20.5 | 20 % |
| se16 | Foxh1 | 27.9 | 30 % |
| se17 | FXR | 16.8 | 35 % |
| se18 | GATA3 | 30.7 | 21 % |
| se19 | GATA3 | 31.2 | 20 % |
| se20 | GATA | 31.7 | 15 % |

**Supplementary Table S4: Homer transcription factor motifs for multi-class classification tasks with 20 classes (MC20):** These motifs are used for MC20 grammars without interactions, with interactions, with interactions with order constraints, and with interactions with spacing constraints. The columns (from left to right) contain the seqgra-internal sequence element ID, the motif name (name of the transcription factor or complex), the minimum KL divergence between the motif and the other 19 motifs in this grammar, and the maximum adjusted empirical similarity score (ESS) between the motif and the other 19 motif in this grammar.

| ID | Motif | min $D_{\mathrm{KL}}(\cdot,\cdot)$ | max ESS$(\cdot,\cdot)$ |
|---|---|---|---|
| se1 | FOXA1:AR | 32.7 | 15 % |
| se2 | Bcl11a | 12 | 45 % |
| se3 | Brachyury | 18.3 | 39 % |
| se4 | CEBP:CEBP | 23.0 | 18 % |
| se5 | Chop | 15.6 | 35 % |
| se6 | CHR | 17.6 | 26 % |
| se7 | CTCF-SatelliteElement | 31.1 | 9 % |
| se8 | Mouse Recombination Hotspot | 24.2 | 10 % |
| se9 | RAR:RXR | 22.3 | 28 % |
| se10 | DUX | 24.1 | 23 % |
| se11 | E2F7 | 20.1 | 15 % |
| se12 | EBNA1 | 35.0 | 12 % |
| se13 | ERE | 16.8 | 20 % |
| se14 | ETS:E-box | 26.9 | 16 % |
| se15 | EWS:ERG-fusion | 17.9 | 18 % |
| se16 | Foxh1 | 20.7 | 36 % |
| se17 | FXR | 13.2 | 38 % |
| se18 | GATA3 | 28.1 | 18 % |
| se19 | GATA3 | 31.2 | 20 % |
| se20 | GATA | 27.4 | 20 % |
| se21 | GATA:SCL | 19.5 | 14 % |
| se22 | Gfi1b | 23.1 | 14 % |
| se23 | GRHL2 | 21.2 | 19 % |
| se24 | Hand2 | 15.2 | 30 % |
| se25 | HINFP | 25.6 | 23 % |
| se26 | HOXB13 | 25.6 | 14 % |
| se27 | LRF | 23.2 | 20 % |
| se28 | LXRE | 20.0 | 28 % |
| se29 | NF1:FOXA1 | 29.1 | 13 % |
| se30 | NFAT:AP1 | 19.1 | 17 % |
| se31 | NFAT | 17.2 | 20 % |
| se32 | NFY | 19.8 | 25 % |
| se33 | Nur77 | 27.6 | 18 % |
| se34 | Oct2 | 17.1 | 29 % |
| se35 | Oct4:Sox17 | 22.6 | 15 % |
| se36 | OCT4-SOX2-TCF-NANOG | 17.9 | 14 % |
| se37 | p53 | 27.3 | 10 % |
| se38 | PAX3:FKHR-fusion | 19.7 | 17 % |
| se39 | PAX5 | 20.8 | 13 % |
| se40 | PAX6 | 23.6 | 14 % |
| se41 | Pax7 | 21.7 | 23 % |
| se42 | Pax7 | 20.7 | 25 % |
| se43 | Pax8 | 25.4 | 18 % |

*Continued on next page*

Supplementary Table S5 – *Continued from previous page*

| ID | Motif | min $D_{\mathrm{KL}}(\cdot,\cdot)$ | max ESS$(\cdot,\cdot)$ |
|---|---|---|---|
| se44 | Pitx1:Ebox | 14.1 | 33 % |
| se45 | PRDM10 | 14.5 | 33 % |
| se46 | PRDM14 | 18.4 | 17 % |
| se47 | PRDM15 | 23.0 | 18 % |
| se48 | PRDM9 | 17.9 | 34 % |
| se49 | PSE | 15.0 | 44 % |
| se50 | RBPJ:Ebox | 17.8 | 24 % |

**Supplementary Table S5: Homer transcription factor motifs for multi-class classification tasks with 50 classes (MC50):** These motifs are used for MC50 grammars without interactions, with interactions, with interactions with order constraints, and with interactions with spacing constraints. The columns (from left to right) contain the seqgra-internal sequence element ID, the motif name (name of the transcription factor or complex), the minimum KL divergence between the motif and the other 49 motifs in this grammar, and the maximum adjusted empirical similarity score (ESS) between the motif and the other 49 motif in this grammar.
